# Default Mode Network activation at task switches reflects mental task-set structure

**DOI:** 10.1101/2024.07.08.602546

**Authors:** Ashley X. Zhou, John Duncan, Daniel J. Mitchell

**Affiliations:** MRC Cognition and Brain Sciences Unit, University of Cambridge

**Keywords:** cognition, default mode network, functional MRI, MVPA, switching

## Abstract

Recent findings challenge traditional views of the Default Mode Network (DMN) as purely task-negative or self-oriented, showing increased DMN activity during demanding switches between externally-focused tasks (Crittenden et al., 2015; Smith et al., 2018; Zhou et al., 2024). However, it is unclear what modulates the DMN at switches, with transitions within a stimulus domain activating DMN regions in some studies but not others. Differences in the number of tasks suggest that complexity or structure of the set of tasks may be important. In this fMRI study, we examined whether the DMN’s response to task switches depends on the complexity of the active set of tasks, manipulated by the number of tasks in a run, or abstract task groupings based on instructional order. Core DMN activation at task switches was unaffected by the number of currently relevant tasks. Instead, it depended on the order in which groups of tasks had been learnt. Multivariate decoding revealed that Core DMN hierarchically represents individual tasks, task domains, and higher-order task groupings based on instruction order. We suggest that, as the complexity of instructions increases, rules are increasingly organized into higher-level chunks, and Core DMN activity is highest at switches between chunks.

## 1 Introduction

The default mode network (DMN) is a set of regions in the human brain that show elevated activity at rest, contrasting with attenuation during tasks that demand externally-focused attention (Buckner et al., 2008). Prominent views suggest that the DMN may be engaged during construction of internal, especially autobiographical, mental models, or during exploratory monitoring of the environment in the absence of attentional focus. This pattern of responding typically contrasts with networks such as the multiple-demand network (Duncan, 2010) which are active in attentionally demanding situations, and often anti-correlate with the DMN (Fox et al., 2005). Subsequent research showed that the set of default mode regions can be further separated into “Core,” “MTL” and “dMPFC” subnetworks (Andrews-Hanna et al., 2010, 2014; Wen, Mitchell, et al., 2020), which have partially specialised roles in various internally-focused cognitive processes such as self-referential processing (Kelley et al., 2002), mind-wandering (Christoff et al., 2009), autobiographical memory and imagery (Addis et al., 2007; Schacter et al., 2007), and social cognition (Mars et al., 2012).

Complicating this broadly consistent picture, recent findings of increased DMN activity, especially of its Core subnetwork, during difficult and externally driven task switches challenge simple views of DMN as a purely ‘task-negative’, or internally focused network (Crittenden et al., 2015; Kurtin et al., 2023; Smith et al., 2018; Zhou et al., 2024). However, studies requiring people to alternate between two, often similar, tasks, typically activate multiple-demand-like regions, rather than the DMN (Braver et al., 2003; Kim et al., 2012; Monsell, 2003). The studies reporting DMN activity during task-switching have instead used larger, often hierarchically organised, sets of tasks. Two of these studies (Crittenden et al., 2015; Smith et al., 2018) used six tasks, grouped into three domains, and found that Core and MTL subnetworks of the DMN responded to between-domain switches, but not to within-domain switches. More recently, we investigated DMN activity as people switched between four tasks, grouped into two domains (Zhou et al., 2024). Although we replicated a DMN task switch effect, showing increased Core DMN activity at task switches versus task repeats, we found comparable DMN activity for all types of task transitions, including within-domain switches. The reduced number of domains in the latest study suggests that the number of tasks may influence whether the DMN is recruited by task switches, and further raises the question of the role of the DMN in externally-driven cognitive transitions.

To investigate whether the DMN task-switch effect depends on the complexity or organisation of the set of tasks, we therefore designed a paradigm that preserves the task-switching trial structure, while manipulating the set of tasks in two critical ways. Firstly, since the number of tasks may constitute a threshold for the involvement of DMN, we compared runs in which participants had to switch between eight different tasks (from four stimulus domains) with runs in which participants were only asked to perform four of these tasks (from two stimulus domains). Secondly, to test whether the DMN might also be sensitive to higher-order, hierarchical mental organisation of the set of tasks, we had participants learn and practise the tasks in two groups of four, allowing us to examine switches between learnt groups, and effects of learning order.

We hypothesised, firstly, that if the Core DMN task-switch response depends on the number of tasks to be performed, then within-domain and between-domain switches would differ more in runs with four domains (replicating Crittenden et al., 2015; Smith et al., 2018) than in runs with two domains (replicating Zhou et al., 2024). Secondly, we hypothesised that if the Core DMN is sensitive to higher-order task structure, then this could be reflected in a greater univariate response for between-group switches compared to within-group switches, a different response to tasks learned later compared to tasks learned first, and a multivariate response pattern that distinguishes tasks from different learning groups better than those from the same leaning group.

## 2 Methods

### 2.1 Participants

38 native English speakers between the age of 18-45 (64.7% female), and with normal or corrected-to-normal vision and no history of neurological or psychiatric disorders, were recruited from the Cognition and Brain Sciences Unit’s healthy participant panel. Two participants were excluded from analysis due to technical errors in presenting stimuli during the experiment and saving data. Participants all provided informed consent and were given monetary compensation for their time. The experiment was conducted in accordance with ethical approval granted by the Cambridge Psychology Research Ethics Committee (CPREC).

Estimated effect sizes were not available based on existing literature, so the choice of sample size assumed a medium standardized effect size of Cohen’s d = 0.5 for the key hypothesis that Core DMN’s within-domain and between-domain switch responses would differ more in four-domain than two-domain runs. A two-tailed t-test with alpha of 0.05 and power of 0.8 yielded a minimum required sample size of 34. Secondary hypotheses are orthogonal, and the same logic and sample size apply. 38 participants were recruited on the expectation that some may have needed to be excluded.

### 2.2 Task Design

The experimental paradigm, adapted from Crittenden et al. (2015) and Smith (2018) and depicted in Figure 1, comprised eight different colour-task pairs. The colours red and green framed an object, and cued ‘Is it living?’ and ‘Does it fit into a shoebox?’ respectively. The colours blue and yellow framed an incomplete word and respectively cued the lexical tasks ‘Does I fit in to make a word?’ and ‘Does A fit in to make a word?’ The pink and cyan colours framed a face and indicated ‘Is this person female?’ and ‘Is this person young?’ Finally, the orange and purple colours framed two shapes and asked ‘Are these the same shape?’ and ‘Are these the same height?’

**Figure 1.**
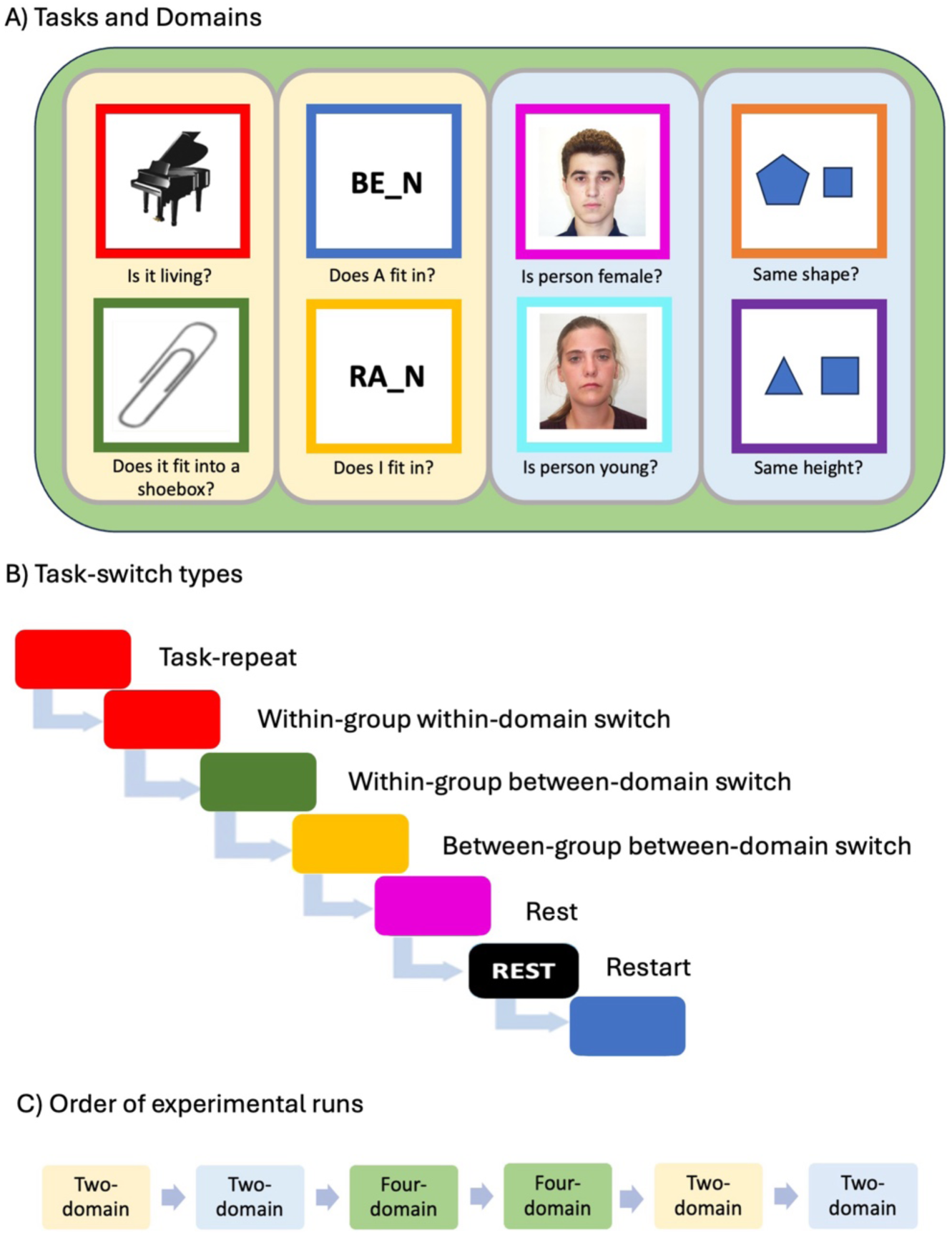
Experimental paradigm. Panel A shows the eight tasks used in the experiment, each cued by a differently coloured square frame. To illustrate the hierarchical structure of the total set of tasks, tasks are encircled in their corresponding domains (semantic, lexical, faces, shapes) and a sample pairing of training groups is indicated by light yellow and light blue backgrounds. B) shows a sample sequence of tasks that generates the relevant task-switch conditions (e.g. task repeat, within-domain switch, between-domain switch). C) shows the sequence of runs in the scanner, which consists of four runs with tasks from only two domains (from a single learning group), and two runs with tasks from all four domains.

On each trial, a coloured frame (4.36 x 4.36 degrees of visual angle) appeared in the middle of the screen, surrounding a simultaneously presented stimulus and cueing the task to be performed. Rest trials were indicated by a black frame, with the participant not required to respond. Each trial ended when the participant responded, or, for rest trials, after the mean reaction time of all preceding trials. The next trial followed after an inter-trial interval of 1.5 seconds, with a central fixation cross.

Participants learnt these eight different task types, two in each stimulus domain, in two practice sessions before entering the scanner. Each session included learning and practising the colour-cue and task rules for two of the four domains. 11 participants learnt the word and face tasks first, followed by the object and shapes tasks; 12 participants learnt the object and shapes task first, followed by the word and face tasks; 13 participants learnt the object and word tasks first, followed by the faces and shapes tasks. During this pre-scan training, participants were first shown the colours that corresponded to each rule, listened to the experimenter describe the rules, and read the instruction sheet until they were confident that they had learnt the pairings. Then, participants practised the tasks cued by the coloured frames, one colour/rule pair at a time, for ten repeats. Participants were then asked to repeat out loud which question each colour cued and to complete a block of 30 trials with mixed cues and a random order of trials. Participants were allowed to proceed to the testing phase if their performance on the practice block exceeded 80%, otherwise they attempted another practice block. After learning the first four tasks in this way, the participant moved on to a similar practice session for the second set of four.

In the main experiment, across each run, we pseudorandomised the sequence in which the tasks were presented, to create different switch conditions (Figure 1B). For two-domain runs, switch types were task repeat, within-group-within-domain switch, within-group-between-domain switch, task-to-rest switch, and rest-to-task switch (“restart”). In four-domain runs, there were additionally between-group-between-domain switches. Within each run, equal numbers of trials per switch condition were counterbalanced across task types. Participants completed four runs with tasks from only two domains, in the same pairs they were learnt, and two runs with trials intermixed from all four domains (Figure 1C). Each two-domain run had 81 trials, 16 for each of five types of task switch (see above), plus a dummy trial with a random task at the start of the run; each four-domain run had 193 trials, with 32 for each of six types of task switch, following an initial dummy trial.

In the first three runs, each task was restricted to be followed by only one of the possible tasks per switch type. For example, if the task on trial N was “Is it living?” (Figure 1A), then for trial N+1, a task repeat trial would also be “Is it living?”, a within-group-within-domain switch trial would be “Does it fit into a shoebox?”, a within-group-between-domain switch trial would always be the same one of the two possibilities, e.g. “Does A fit in?”, and in a four domain run, a between-group-between-domain switch trial would again always be only one of the four possibilities (e.g. “Is person a female?”). In this way the probability of encountering a particular task transition was fixed across trial types. In the second three runs of the experiment, this constraint was removed, so that, for a given trial type, any of the possible next tasks could occur. fMRI findings were very similar for the two halves of the session, so all results from the two were averaged.

### 2.3 MRI acquisition and regions of interest (ROIs)

Data were acquired on a 3T Siemens Prisma MRI scanner fitted with a 32-channel coil. A structural T1-weighted structural scan was acquired using an MPRAGE sequence (TR 2.4s, TE 2.2ms, flip angle 8°, voxel size 0.8 × 0.8 × 0.8 mm). Task runs used T2*-weighted Echo-Planar Imaging (multiband acquisition factor 2, TR 1.2 s, TE 30 ms, flip angle 67°, voxel size 3 × 3 x 3 mm).

Pre-processing involved spatial realignment, slice-time correction, co-registration, and normalization to the MNI template brain (using SPM12 and AutomaticAnalysis; Cusack et al., 2015). No spatial smoothing was used for multivariate or region of interest (ROI) analyses.

Within the DMN, we focused on the Core subnetwork because this has shown the largest and most consistent response to task switches in previous studies (Crittenden et al., 2015; Zhou et al., 2024). The Core DMN ROI was taken from Wen, Mitchell et al (2020), based on local voxel clusters from the Yeo et al (2011) 17-network parcellation that contain the Core subnetwork’s coordinates provided in Andrews-Hanna et al. (2010). (See Wen, Mitchell et al. 2020 for further details.)

### 2.4 Univariate fMRI Analysis

For each participant, a general linear model was created, with regressors for each combination of switch condition (task repeat, within-group-within-domain switch, within-group-between-domain switch, between-group-between-domain switch, restart, rest), number of task domains in the run (two or four), task expectancy (constrained transitions, all transitions), and learning group (first or second). Regressors were created by convolving the response periods (from stimulus onset to trial end), of all trials per condition, with the canonical hemodynamic response function. Six rigid-body realignment parameters were included as non-convolved regressors per run, along with run means. The model and data were temporally high-pass filtered using the default cut-off period of 128 s. Analysis focused on the mean signal across the Core DMN ROI, and compared switch conditions using repeated measures ANOVAs and paired t-tests. Frequentist statistics were supplemented with default Bayes factors based on F statistics for ANOVAs (Faulkenberry & Brennan, 2023) and t statistics for t-tests (Rouder et al., 2009).

### 2.5 Multivariate fMRI Analysis

Neural discrimination of each pair of tasks was examined by training a linear support vector machine (SVM) classifier (LIBLINEAR, Fan et al., 2008) to classify tasks, for each ROI, and each participant, using The Decoding Toolbox (Hebart et al., 2015). Cross-validation was performed across runs, testing separately on held-out four-domain and two-domain runs, and training on all possible combinations of remaining runs for which classes and domain number were balanced. Classification performance was quantified as the signed distance of test patterns from the decision boundary. This measure was chosen because it gives a continuous, unbounded, and unbiased estimate of representational distance, while being insensitive to differences in pattern variance between conditions (Nili et al., 2020). Classifier performance was compared to chance using two-tailed t-tests, and compared across switch conditions using repeated measures ANOVA and paired two-tailed t-tests.

## 3 Results

### 3.1 Behavioural results - Switch costs did not depend on the number of currently relevant tasks

Mean accuracy was consistently high, at 96% across all trials. Mean reaction time (RT) for correct trials (Figure 2) was highly similar in two-domain runs (1.64 s), and in four-domain runs (1.75 s). Switch costs were evident, where average reaction times for task switches (combined across types) were significantly higher than task-repeat trials in both the two-domain runs (t_35_ = 9.85, p<0.01, BF_10_ =7.33 x 10^8^) and the four-domain runs (t_35_ = 9.92, p<0.01, BF_10_ =8.72 x 10^8^).

**Figure 2.**
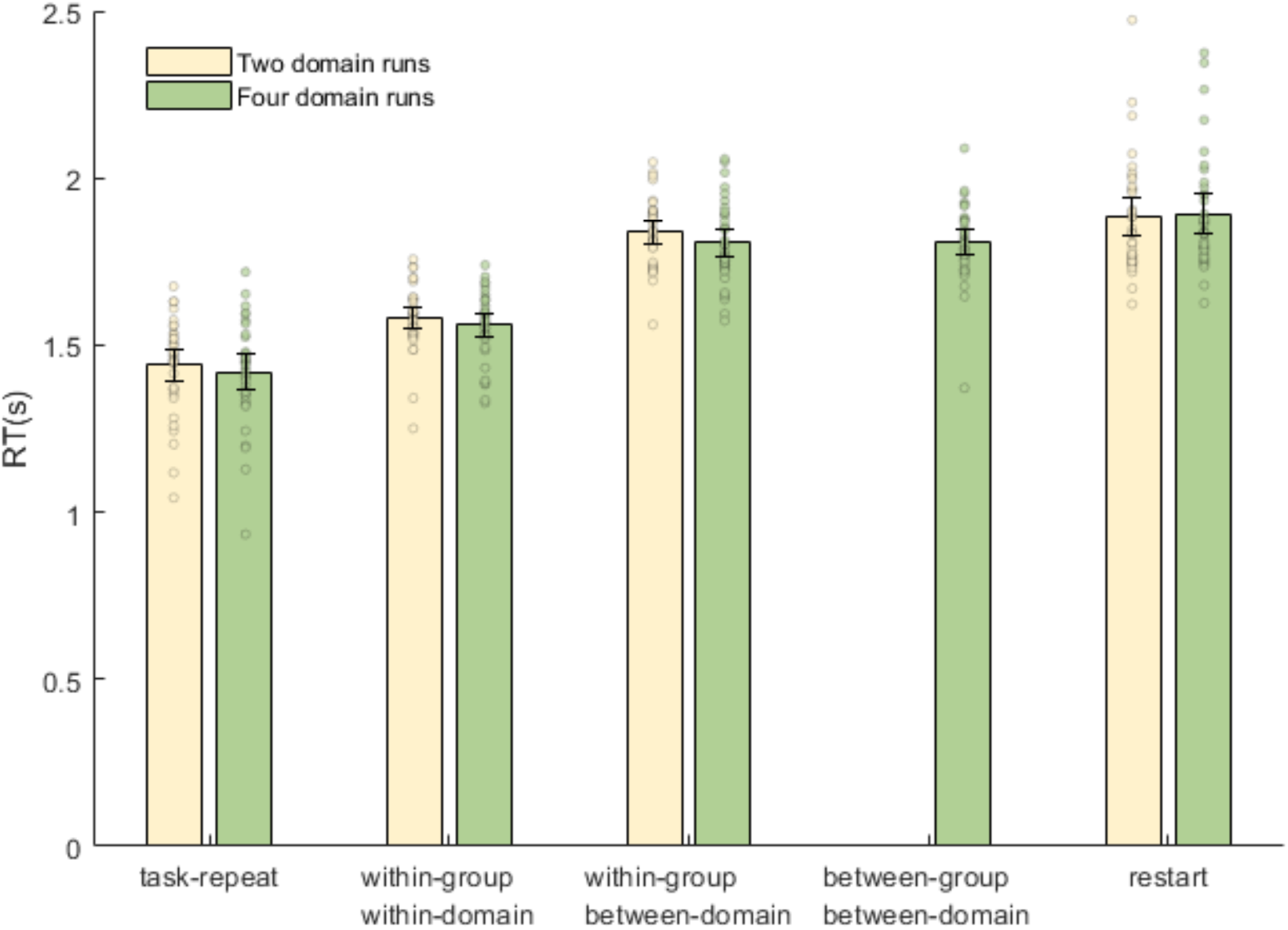
Reaction times for each task transition type in two- and four-domain runs. Switch costs were observed, but surprisingly RTs were not slower in four-domain runs (green) than two-domain runs (cream). Error bars indicate within-subject 95% confidence intervals. Dots show data for individual participants, after removal of between-participant variance (Loftus & Masson, 1994).

Reaction times were examined further using a three-way, repeated-measures ANOVA with factors of switch type (task-repeat, within-group-within-domain, within-group-between-domain, and restart), number of domains present in the current run (two or four), and the learning group to which the task belongs (first or second). Switch type showed a significant main effect (F_(3,105)_ =91.13, p<0.01, BF_10_=8.71 x 10^25^), while other main effects and interactions were non-significant (F<1.33, p>0.26, BF_10_<0.39).

Post hoc, paired t-tests across switch types found significantly higher reaction times for within-group-within-domain switch trials compared to task repeat trials (t_35_=7.75, p<0.01, BF_10_ =2.93x10^6^), and significantly higher reaction times for within-group-between-domain switch trials than within-group-within-domain switch trials (t_35_=11.40, p<0.01, BF_10_ =3.10 x 10^10^). While restart trials had significantly higher reaction times compared to within-group-within-domain switch trials (t_35_=8.23, p<0.01, BF_10_ =1.16 x 10^7^), no significant difference was observed between restart and within-group-between-domain switch trials (t_35_=1.97, p=0.06, BF_10_ =1.00).

To investigate effects of task switches between tasks from different learning groups, a two-way, repeated-measures ANOVA using only trials from the four-domain runs assessed factors of switch type (within-group-between-domain, between-group-between-domain) and learning group. There was no significant effect of switch type, learning group, or interaction (F_(1,35)_ <0.70, p>0.41, BF_10_<0.30).

### 3.2 Univariate fMRI Analysis

#### 3.2.1 Simple switch effect replicated in Core DMN

In the Core DMN ROI, we first examined the basic switch effect, contrasting against task-repeat trials, and averaging across all other factors (Figure 3A). Replicating previous findings of the DMN switch effect (Crittenden et al., 2015; Kurtin et al., 2023; Smith et al., 2018; Zhou et al., 2024), a t-test confirmed a significant increase in Core DMN activity at task switches (averaged across switch types) compared to task repeats (t_35_ =4.46, p<0.01, BF_10_=301). A t-test also found greater Core DMN activity for within-group-between-domain switches compared to within-group-within-domain switch trials (t_35_=4.43, p<0.01, BF_10_=280).

**Figure 3.**
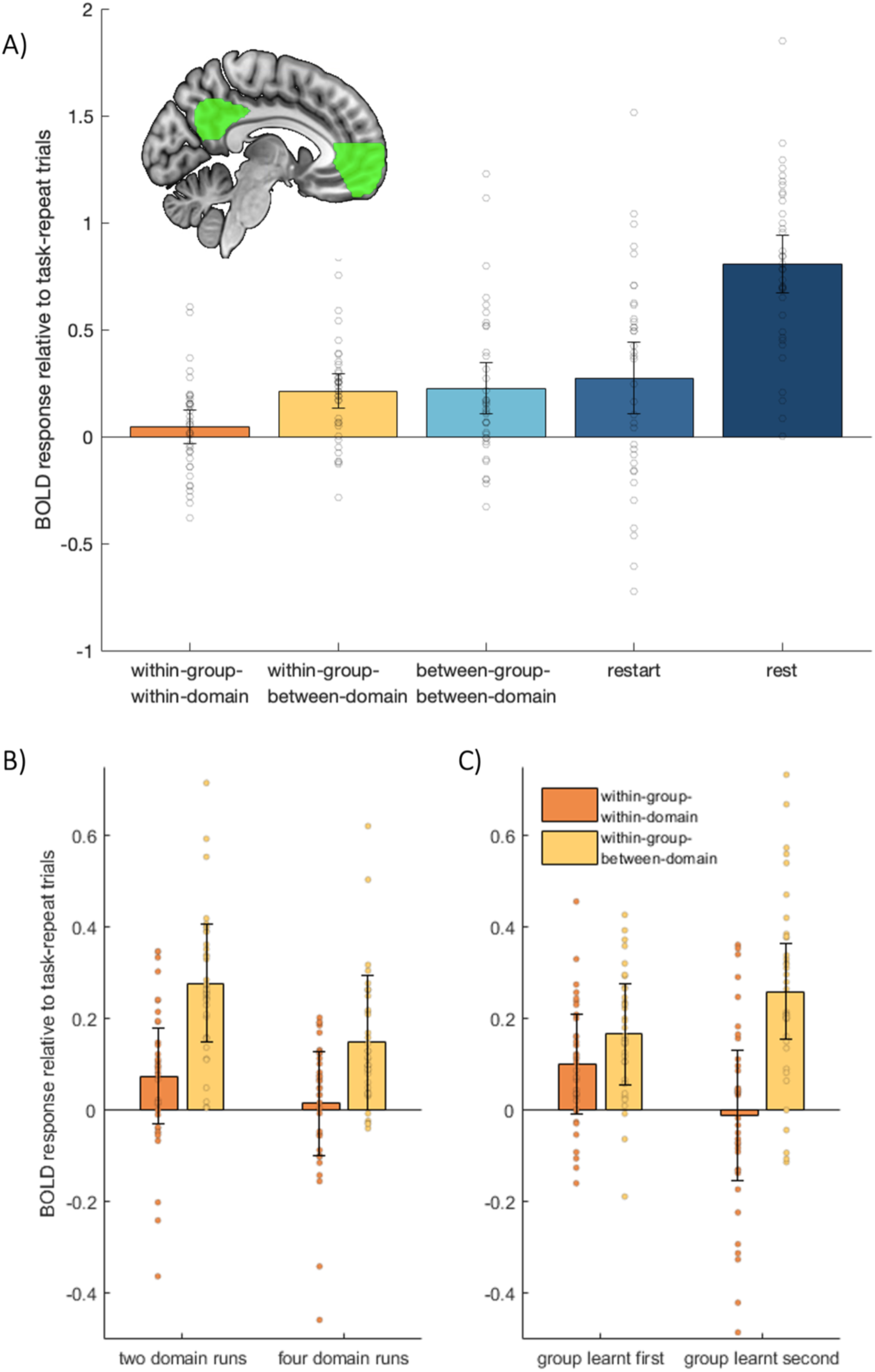
Core DMN BOLD response. (A) shows the Core DMN’s BOLD response for each task transition type, relative to task-repeat. The difference between within-group-within-domain and within-group-between-domain transitions is further broken down according to runs with tasks from two or four domains (B), and whether the tasks belonged to the first or second learning group (C). (D) shows the Core DMN ROI. Error bars indicate between-subject 95% confidence intervals for the contrast of each condition against the task repeat baseline. Dots show data for individual participants, after removal of between-participant variance (Loftus & Masson, 1994).

Next, we tested whether responses to between-group-between-domain transitions were greater than for within-group-between-domain transitions in four-domain runs, as evidence for a higher-order hierarchy in the switch effect. However, matching the behavioural results, transitions between different learning groups did not significantly differ from transitions between domains within a group (t_35_=0.30, p=0.76, BF_10_=0.19).

Also in line with behaviour, the response at transitions from rest back to task (restart) was significantly greater than at within-group-within-domain switches (t_35_=2.73, p=0.01, BF_10_=4.24), but not significantly different from within-group-between-domain switches (t_35_=0.85, p=0.40, BF_10_=0.25) or between-group-between-domain switches (t_35_=0.57, p=0.57, BF_10_=0.21).

Finally, we compared the average of all task-switch trials to rest trials. This confirmed that the Core DMN ROI was significantly deactivated by the task trials compared to the rest trials, as expected for a DMN region (t_35_=9.73, p<0.01, BF_10_=5.38x10^8^).

#### 3.2.2 Core DMN switch effects do not depend on complexity of the current set of tasks, but do depend on the order in which tasks were learnt

We next examined the effect of each hypothesised modulator of the task-switch effect, using repeated measures ANOVAs, in which transition types of interest (within-group-within-domain and within-group-between-domain) were crossed with the two modulating factors of current task-set complexity (two domains, four domains) and instructed order (learnt first, learnt second). Since some studies (Crittenden et al., 2015; Smith et al., 2018) had found a difference between within-domain and between-domain switches, but our most recent study had not (Zhou et al., 2024), we first focused on these two transition types, and were specifically interested here in their interaction with each potential modulating factor. To ensure that effects of learning order did not depend on differential responses to particular stimulus domains, these ANOVAs also included a between-participant factor of which two domains were learnt first (words and faces, objects and shapes, words and objects).

Contrary to prediction, there was neither a significant main effect of domain number (F_(1,33)_ =0.09, p=0.77, BF_10_=0.23), nor an interaction with transition type (F_(1,33)_ =1.14, p=0.29, BF_10_=0.37), suggesting that increasing the complexity of the current set of tasks did not increase the difference between switch types (Figure 3B). Interestingly, however, there was an interaction of switch type with the learned order of the domains (Figure 3C), with higher activity for between-domain switches of later-learned domains (F_(1,33)_ =5.23, p<0.05, BF_10_=2.03). There was no main effect of learned order (F_(1,33)_ =0.42, p=0.52, BF_10_=0.27). Paired t-tests revealed no significant difference between within-domain and between-domain switch conditions within the first learnt group (t_35_=1.45, p=0.16, BF_10_=0.46), replicating Zhou et al. (2024) but a significant increase in Core DMN activity in between-domain compared to within-domain conditions for the second-learnt group (t_35_=3.92, p<0.01, BF_10_=72.72), replicating Crittenden et al. (2015) and Smith et al. (2018). There was no significant main effect of the between-participant factor of domain-pairing (F_(2,33)_ =1.54, p=0.23, BF_10_=0.24), nor interactions with factors of transition types, domain number or learned order (F_(2,33)_ <0.91, p>0.41, BF_10_<0.14), indicating that the effect of learning order generalised across the particular group of tasks learnt first.

Next, to confirm that the Core DMN response to between-domain switches is similar whether within or between groups, even when accounting for learning order, we ran a second ANOVA using only between-domain switches from four-domain runs and crossing transition type (within group, between group) with instructed order (learnt first, learnt second). The types of domains learnt first (words and faces, objects and shapes, words and objects) were again added as a between-subject factor. There was a main effect of instructed order (F_(1,33)_ =5.80, p=0.02, BF_10_=2.53), with activity higher for later-learned tasks. Neither the main effect of transition type, nor interaction, were significant (F_(1,33)_ <3.87, p>0.06, BF_10_<1.18). Again, there was no significant effect of the between-subject factor of type of domains learnt first (F_(2,33)_=0.86, p=0.43, BF_10_=0.13), nor any interaction with transition type or instructed order (F_(2,33)_<1.47, p>0.24, BF_10_<0.22).

### 3.3 Multivariate fMRI analysis

#### 3.3.1 Core DMN response patterns hierarchically discriminate tasks, domains, and learning groups

To assess whether the distinctions between tasks, domains, and learning groups were represented within the Core DMN, we performed multivariate pattern analysis to distinguish every pair of tasks. In Figure 4, the average task-pair discrimination strength is plotted, separately for task pairs that come from the same domain, from different domains within the same learning group, or from different learning groups, and separately for two-domain and four-domain runs. A three-by-two ANOVA with factors of task-pair relationship and domain number found significant main effects of domain number (F_(1,35)_ =9.03, p<0.01, BF_10_=8.53), task-pair relationship (F_(2,70)_ =194.33, p<0.01, BF_10_=6.48x10^25^), and a significant interaction (F_(2,70)_ =11.10, p<0.01, BF_10_=294).

**Figure 4.**
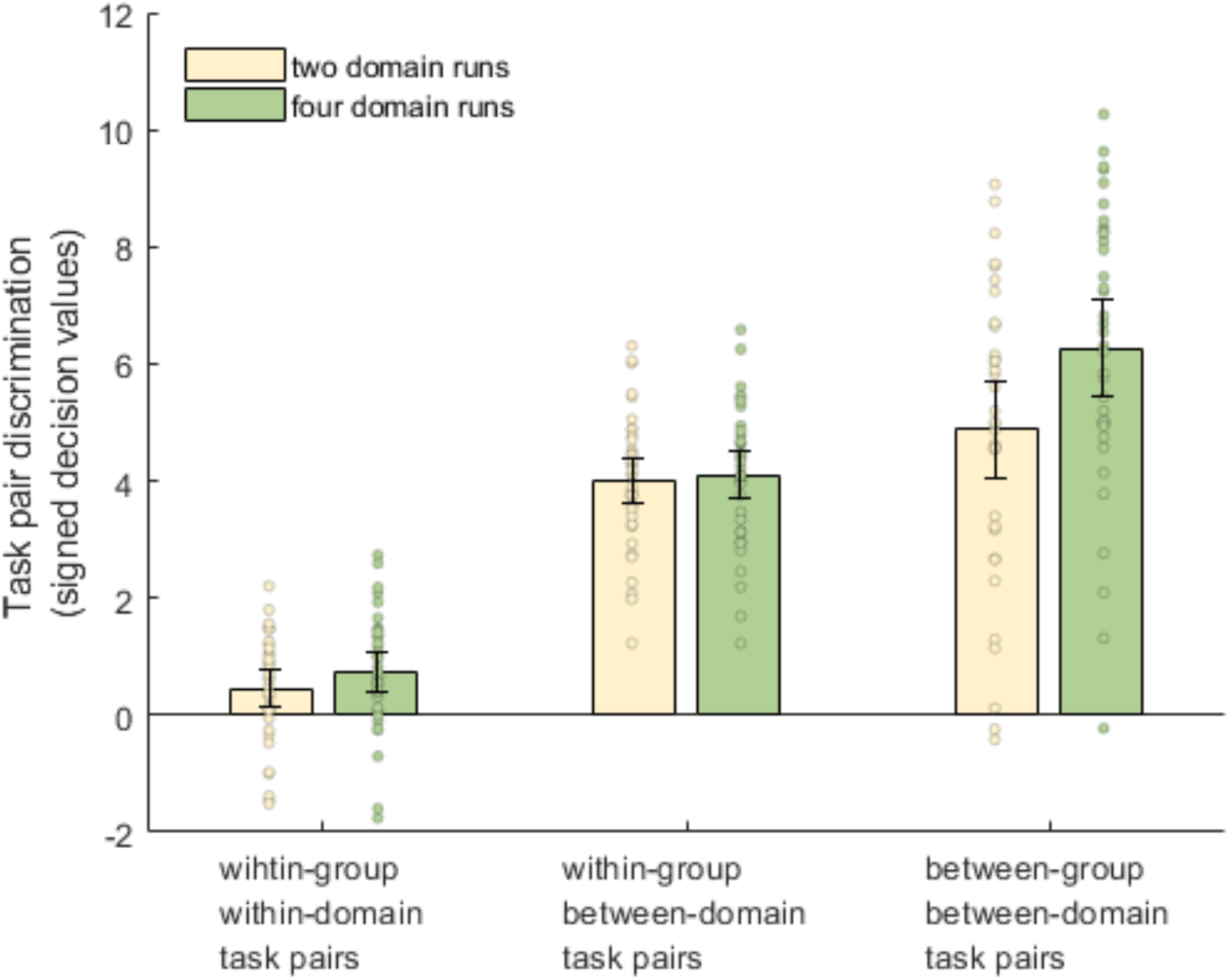
Classification performance for task pairs. Core DMN classification accuracies for task pairs split by domain number and the relationship of the task pair. Error bars indicate 95% between-subject confidence intervals. Dots show data for individual participants, after removal of between-participant variance (Loftus & Masson, 1994)

*Post hoc* two-tailed t-tests confirmed stronger discrimination of within-group-between-domain task pairs than within-group-within-domain task pairs, for two domain runs (t_35_=17.36, p<0.01, BF_10_=5.37 x 10^15^), and also for four domain runs (t_35_=15.53, p<0.01, BF_10_=1.91 x 10^14^), consistent with previous results (Crittenden et al., 2015; Smith et al., 2018) and as expected based on perceptual differences between stimulus domains. Extending this result, two-tailed t-tests revealed stronger discrimination of between-group-between-domain task pairs than within-group-between-domain pairs, both for two-domain runs (t_35_=2.73, p<0.01, BF_10_=4.28) and also for four-domain runs (t_35_= 8.29, p<0.01, BF_10_=1.25 x 10^7^). This implies that the Core DMN also represents the abstract task structure imposed by the learning groups. Interestingly, a two tailed t-test showed that between-group task pairs were also discriminated better in four-domain runs (when tasks from both groups were performed), than in two-domain runs (t_35_= 3.33, p<0.01, BF_10_ = 16.66).

Because domains were assigned to learning groups such that the face tasks were never learnt in the same group as the object or word tasks, it was important to confirm that greater discrimination of between-group task pairs could not be due to greater dissimilarity of these particular task combinations. Therefore, the comparison of between-group-between-domain pairs versus within-group-between-domain pairs was repeated, but excluding those task pairs that were always between-group, and adding the between-participant factor of first-learnt domains (words and faces, objects and shapes, words and objects). The within-participant factor of domain number was also included. This three-way mixed ANOVA showed no significance for the between-participant factor or any interaction with it (F <1.15, p>0.33, BF_10_<0.17). The significant main effects of both domain number (F _(1,33)_ =7.05, p=0.01, BF_10_=4.08), and task-pair type (F _(2,66)_ =191.49, p<0.01, BF_10_=5.22x10^24^), were retained, along with their interaction (F _(2,66)_ =8.82, p<0.01, BF_10_=53.62). Therefore, the greater neural discrimination of tasks learnt in different groups generalised across the particular identity of these tasks.

Finally, we assessed whether discrimination of within-group-within-domain or within-group-between-domain task pairs depended on whether the tasks were from the first or second learning groups. In the first learning group, participants were assigned either the object and word domains, object and shape domains, or word and face domains; thus this analysis was restricted to the latter two subsets, for whom domains were balanced across first and second learning groups. A three-way mixed ANOVA included within-participant factors of task-pair type (within-group-within-domain, within-group-between-domain) and learning group (first, second), plus the between-participant factor of first-learnt domains. Aside from the previously reported effect of task-pair type (F _(1,21)_ =110.59, p<0.01, BF_10_=4.23 x 10^6^), there was no significant effect of learning order (F _(1,21)_ =6.46x10^-5^, p=0.99, BF_10_=0.28), and no significant interaction of task-pair types and learning order (F _(1,21)_ =2.12, p=0.16, BF_10_=0.67). There was also no significant main effect of the between-participant factor of types of domains learnt first (F _(1,21)_ =4.26x10^-3^, p=0.95, BF_10_=0.28) and no significant interaction with task-pair types (F _(1,21)_ =0.35, p=0.56, BF_10_=0.33), although a significant interaction with learning order (F _(1,21)_ =21.08, p,<0.01, BF_10_=147) suggested that task discrimination depended on the identity of the tasks.

## 4. Discussion

### Core DMN switch effects vary with instructional complexity of the overall set of tasks, but not the complexity of current demands

This study set out to investigate factors that affect recruitment of default mode regions at external task switches, and to resolve an inconsistency across studies, whereby equal Core DMN response to within-domain and between-domain task switches has been observed with four tasks organised into two domains (Zhou et al., 2024), but preferential response to between-domain switches has been observed with six tasks organised into three domains (Crittenden et al., 2015; Smith et al., 2018). Here, we report imaging results that replicate higher activation of Core DMN at task switches overall, along with a tendency for higher activity on between-domain shifts modulated by overall complexity. Specifically, we found no evidence that the switch effect differed for four-domain runs compared to two-domain runs. Instead, we found a significant interaction between learning group and task switch type. For the tasks learnt first, Core DMN activity was similar for within- and between-domain switches, replicating Zhou et al (2024). For the tasks learnt second, activity was greater for between-domain switches, replicating Crittenden et al. (2015) and Smith et al. (2018). These results suggest that the Core DMN is sensitive to a mental task structure established during task learning. With accumulating complexity of the total set of tasks, the later tasks become increasingly chunked by domain, such that Core DMN becomes preferentially sensitive to coarser, between-domain task transitions.

The insensitivity to the complexity of the current set of tasks is also reflected in the behavioural results. Reaction times did not differ between two-domain and four-domain runs, and so did not depend on the number of rules active in the current run. This is a potentially surprising observation, given classic demonstrations that selection from amongst a larger set of alternatives is often more difficult (Criss & Shiffrin, 2004; Hick, 1952; Lewis & Anderson, 1976; Schneider & Anderson, 2011). In our data, reaction time is insensitive to the number of alternatives in the active set of tasks, depending only on the relationship between the current and preceding task. In fact, this result was predicted by Oberauer in his theory of procedural working memory (2009), and has now been found several times (Kessler & Meiran, 2010; Souza et al., 2012; van ’t Wout et al., 2015). As cognitive load increases, we predict that task information contributing to the overall mental task model is organised into chunks, such that reaction time on a given trial will depend on the complexity of the chunk to which the task belongs, more than the complexity of other chunks.

Insensitivity to the complexity of the current set of tasks, but dependence of the Core DMN response on the overall body of instructions, is reminiscent of the phenomenon of ‘goal neglect.’ Goal neglect describes a phenomenon whereby a participant fails to implement a task rule, despite the rule being understood, remembered, and executable after sufficient prompting (Duncan et al., 1996). Echoing the present results, neglect of a task rule within a complex task structure is insensitive to the number of rules active in any one block, but depends instead on the number of rules participants are instructed to remember at the beginning of all the blocks (Bhandari & Duncan, 2014; Duncan et al., 2008).

The Core DMN’s preferential response to between-domain, relative to within-domain, task switches also resembles this ‘goal-neglect’ pattern. While this experiment finds no effect of complexity of the current set of tasks, comparing the results of Zhou et al. (Zhou et al., 2024; four instructed tasks and no preferential between-domain response) to this and prior experiments (Crittenden et al., 2015; Smith et al., 2018; six or eight instructed tasks and a preferential between-domain response) suggests a dependence on total instructional complexity. Another suggestive link to the phenomenon of goal neglect concerns the observation that rules learned later are more likely to be neglected (Duncan et al., 1996). Here, we find that the Core DMN’s preferential response to between-domain switches emerges for the last-learned group of tasks. These observations suggest that it would be worthwhile for future experiments to examine the relationship between behavioural ‘goal neglect’ and Core DMN involvement more explicitly.

### Core DMN activation patterns reflect a multi-level task hierarchy, but its switch response operates at a single level

The effect of instruction was also seen in stronger multivariate discrimination of task pairs from different learning groups. The results show that Core DMN is sensitive to abstract cognitive structures established during learning, confirming that teaching the rules in two groups created distinct mental representations. It also implies that Core DMN activation patterns reflect a multi-level task hierarchy (Wen, Duncan, et al., 2020), with increasingly distinct representations for task pairs respectively within a domain, across domains, and across learning groups.

It is therefore interesting that this stronger neural discrimination of between-group representations was not accompanied by an increase in Core DMN activity, or by greater behavioural switch costs, at between-group task switches relative to within-group-between-domain switches. Despite multi-level task hierarchies being distinguished in its response pattern, Core DMN activation at task switches appears insensitive to this deeper hierarchy. In turn, this suggests that transition-related DMN activity may only reflect the occurrence of a transition between cognitive chunks, without being scaled by the magnitude of the cognitive or neural pattern difference between these chunks. Of course, it remains possible that different or stronger manipulations of task groupings could reveal task-switch responses with a deeper hierarchy, and it would be useful for future studies to test the generality of the current proposal.

### Possible implications for capacity limits in a cognitive task model

A tentative explanation for the current results is that the later-learned task domains exert strain on the cognitive capacity for representing the overall task model, such that later-learned tasks become chunked more strongly by domain, which then evoke stronger recruitment of the Core DMN at switches between these chunks. Such recruitment of default mode regions could reflect various processes, including detachment from previously relevant chunk of task rules, or activation of the requirements of the newly appropriate chunk.

This proposal is consistent with the observation that activation of Core DMN is only observed at some task switches, because activation will depend on the chunking structure of the overall set of tasks, which in turn depends on the similarity of the tasks, the total complexity of all instructions, and individual capacity limits in maintaining the task model. Integrating results across studies suggests two distinct capacity limits that might be at play. A first capacity limit is the number of tasks or rules that can fit into a “chunk”. The lack of a DMN switch response in traditional two-task designs (Kim et al., 2012), even for tasks belonging to different stimulus domains (Smith et al., 2019), suggests that a single chunk can contain at least two tasks. The emergence of a DMN response when switching amongst four tasks (Zhou et al., 2024) suggests that four tasks exceed the capacity of a single large chunk, so are preferentially represented as four smaller chunks. A second limit is the number of chunks that can be readily maintained. For sets of six or more tasks, the re-emergence of chunking into task pairs (Crittenden et al., 2015; Smith et al., 2018; and the later-learnt domains in the current experiment) suggests that beyond four or five chunks it becomes easier to increase the size of a chunk than to add more chunks. These limits likely depend somewhat on the complexity and confusability of individual rules. This idea relates to previous proposals of multi-level procedural memory (Oberauer 2009), and would require further exploration, perhaps varying the number of tasks/rules per stimulus domain, as well as the number of domains, and correlating with individual differences in working memory capacity (Duncan et al., 2012).

Previous work has shown that frontoparietal multiple-demand regions have a phasic response to instruction presentation, which saturates as further instructions are added, and approaches an asymptote after two to three rules (Dumontheil et al., 2011). Although here we did not measure activation during instructions, this gradually saturating frontoparietal response at instruction could plausibly contribute to establishing a task model with well-segregated (less chunked) component rules. This could explain why later-learned tasks collapse into coarser chunks, as the frontoparietal responsiveness asymptotes. It would be interesting for future studies to examine how responses during instruction relate to the results presented here.

Overall, we document sensitivity of the Core DMN to an intrinsic task hierarchy, suggesting a role in complex cognitive control processes. Results help to reconcile inconsistent observations of DMN activation across task-switch studies, and potentially suggest multiple capacity limits within a mental model of the overall set of tasks. We propose that the Core DMN represents a multi-level task hierarchy, whose chunking structure is determined at instruction, and influences activity at transitions between chunks.

## Data and code availability

Code for experiment stimuli, data analysis and figures presented in this manuscript, along with data for generating the figures, can be found here: https://github.com/ashleyxzhou/DMN-TaskSwitch

## Author contributions

A.X.Z., D.J.M. and J.D. designed the research. A.X.Z. collected the data. A.X.Z. and D.J.M. analysed the data. A.X.Z. and D.J.M. wrote the paper. A.X.Z. created the figures. A.X.Z, D.J.M. and J.D. revised the paper. D.J.M. and J.D. supervised the work.

## Acknowledgements

For the purpose of open access, the UKRI-funded authors have applied a Creative Commons Attribution (CC BY) licence to any Author Accepted Manuscript arising from this submission.

Ashley X. Zhou, Daniel J. Mitchell and John Duncan were supported by Medical Research Council Intramural Program MC_UU_00030/7. Ashley X. Zhou was supported by a Gates Cambridge Scholarship.

